# Recombinant GDF11 Improves Neurological Recovery in Models of Hemorrhagic Stroke and Traumatic Brain Injury

**DOI:** 10.64898/2026.06.04.730114

**Authors:** Haichen Wang, Ori S. Cohen, Laura Ben Driss, Viviana Cantillana, Yongting Wang, Manisha Sinha, Anish Deshpande, Tyler Daman, Timothy D. Faw, Daniel T. Laskowitz, Anthony Sandrasagra, Richard T. Lee

## Abstract

**Background:** Intracerebral hemorrhage (ICH) and traumatic brain injury (TBI) are leading causes of long-term neurological disability and mortality worldwide, with no approved therapies that promote functional recovery. Growth differentiation factor 11 (GDF11), a circulating TGFβ-family protein, has shown regenerative and neurorestorative potential in models of ischemic stroke.

**Methods:** We evaluated recombinant GDF11 (rGDF11) in mouse models of ICH and TBI. ICH was induced by intrastriatal collagenase injection, and neurological recovery was assessed using Neuroseverity Score (NSS), Rotarod (RR), and CatWalk (CW) analyses up to 28 days post-injury. Histological assessments of vascularization, neuronal density, and microglial/macrophage density were performed 28 days after ICH. For TBI, a closed-head injury model using a pneumatic impactor was employed, and NSS and RR assessments were conducted through 28 days post-injury.

**Results:** rGDF11 treatment significantly improved neurobehavioral performance following ICH, including NSS, RR, and CW parameters (forelimb base of support and average speed). Histological analyses revealed enhanced vascular area and neuronal density, with reduced microglial/macrophage density in rGDF11-treated mice. Following TBI, rGDF11 accelerated functional recovery, improving RR latency by day 6 and NSS by day 28 post-injury.

**Conclusion:** rGDF11 promotes structural and functional recovery after both hemorrhagic and traumatic brain injury in mice. These findings, together with prior evidence in ischemic stroke, support rGDF11 as a promising neurorestorative biologic with broad therapeutic potential for diverse forms of brain injury.

**Graphical Abstract:** 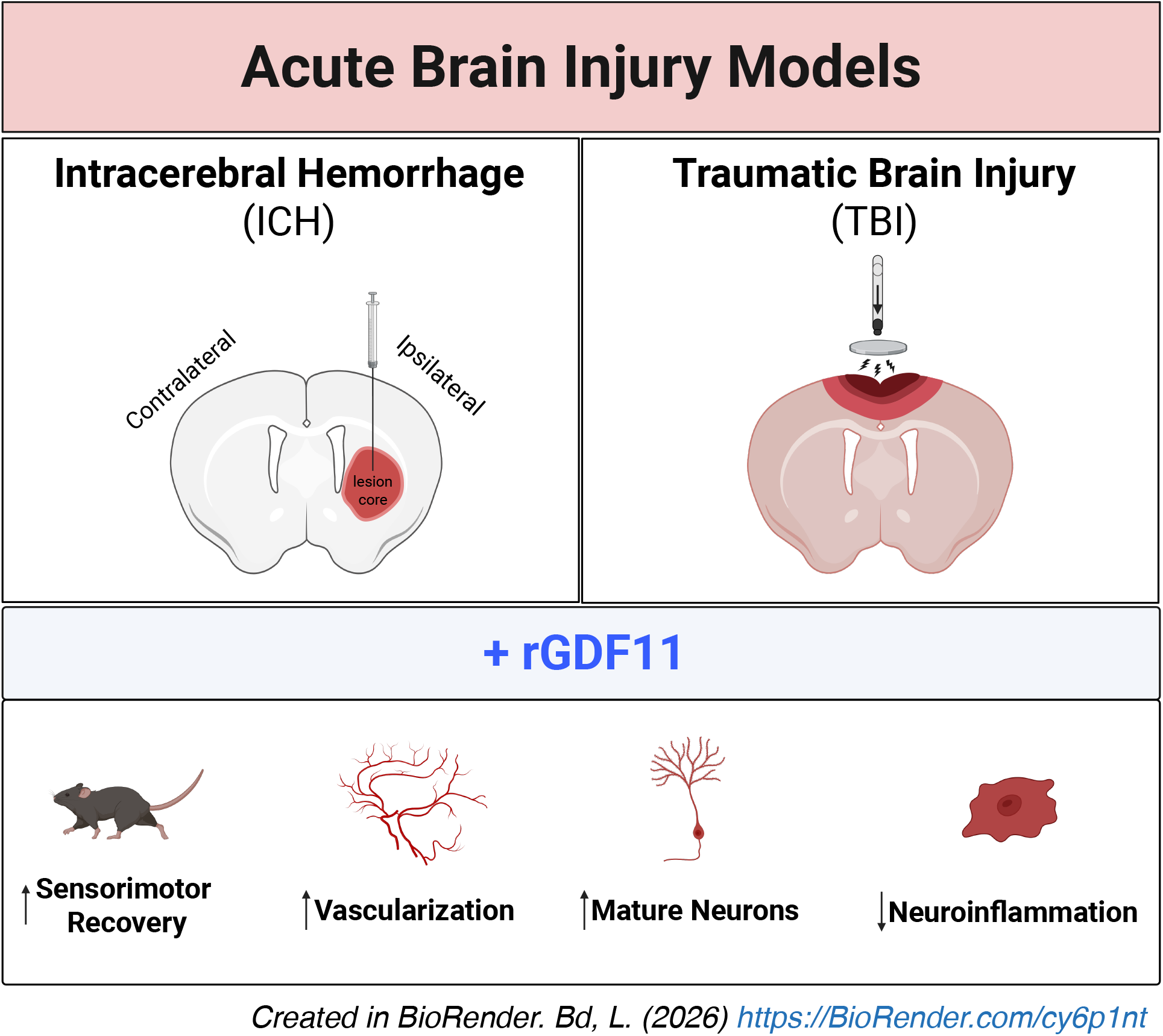

## INTRODUCTION

Intracerebral hemorrhage (ICH) and traumatic brain injury (TBI) are among the leading causes of death and disability worldwide. Despite advances in acute management— including control of blood pressure, seizures, and intracranial pressure—outcomes after ICH remain poor, with 30-day mortality approaching 50%.^1^ Similarly, TBI affects millions annually and frequently results in persistent cognitive and motor deficits, as well as an elevated risk of premature mortality.^2^ Unlike ischemic stroke, where reperfusion therapies are available, there are currently no approved pharmacologic interventions that enhance neurological recovery after ICH or TBI, underscoring the urgent need for neurorestorative treatments that can be administered after injury.

Growth differentiation factor 11 (GDF11), a member of the transforming growth factor-β (TGF-β) superfamily, has emerged as a circulating factor that modulates vascular, neuronal, and inflammatory processes in multiple models of age-related dysfunction.^3-7^ The effects of GDF11 on tissue repair and aging appear to be both concentration- and context-dependent, highlighting the need for comprehensive assessment across biologically relevant disease models. Our recent work demonstrated that systemically delivered recombinant GDF11 (rGDF11) enhances neovascularization, reduces inflammation, promotes neurogenesis, and improves sensorimotor function in a rat model of ischemic stroke.^5^ These findings suggest that rGDF11 acts as a systemic regenerative cue that orchestrates coordinated vascular and neuronal repair after brain injury, providing a rationale for testing its therapeutic potential in other forms of cerebral injury.^5-7^

Here, we evaluated the effects of rGDF11 in mouse models of ICH and TBI using randomized, blinded, and objectively quantified assessments of motor and behavioral recovery. We found that rGDF11 significantly improved neurological function following both injuries, accompanied by evidence of enhanced vascular and neuronal integrity and reduced neuroinflammation. Notably, rGDF11 accelerated recovery within the first week after injury, consistent with its previously observed effects in ischemic stroke.^5^ Together, these findings position rGDF11 as a promising candidate for post-injury neurorepair and motivate further investigation into the mechanisms underlying its reparative actions in the injured brain.

## METHODS

### Data Availability Statement

The data that support the findings of this study are available from the corresponding authors upon reasonable request.

### Protein Production

rGDF11 (recombinant human GDF11 mature domain dimer) was isolated and purified from mammalian cell culture supernatants either from transient expression in HEK Expi293 cells (ThermoFisher Scientific) or in a CHO cell line (CHOZN® GS-/- ZFN-modified, Millipore Sigma) engineered to stably express rGDF11, generated as described previously ^5^ and detailed in the Supplemental Methods.

### Animal Welfare Statement

Male C57BL6J mice (12 – 14 week old, The Jackson Laboratory, Bar Harbor, Maine, USA) were housed in a temperature and humidity controlled environment with a 12 hr light-dark cycle and were allowed free access to food and water. The animals were randomized into experimental groups, and all procedures, assessments, and data analyses were performed in a blinded fashion. All animal studies were approved by Duke University Institutional Animal Care and Use Committee and conducted in accordance with institutional and federal guidelines for animal care. Experiments adhered to ARRIVE Guidelines 2.0 to ensure rigor and reproducibility. ^8^

### Intracerebral Hemorrhage

For ICH, intrastriatal collagenase injection was performed on mice as previously described ^9-11^, with detailed methods provided in the Supplemental Material. rGDF11 or vehicle was administered to male mice by intraperitoneal injection (1 mg/kg) starting 30 minutes after ICH and then every 24 hours for 7 days. Behavioral assessments including Neuroseverity Score (NSS)^9,12,13^ and Rotarod Latency (RR)^9,14^ were conducted pre-ICH and on days 1-7, 14, 21, and 28 post-ICH. CatWalk (CW) assessment was performed pre-ICH, and on day 7 post-injury.^15^

### Traumatic Brain Injury

For TBI, the murine closed head injury model with a pneumatic impactor was employed as described by Laskowitz et al.,^16^ with detailed methods provided in the Supplemental Material. rGDF11 or vehicle was administered to 12–14 week old male C57BL/6J mice by intraperitoneal injection (1 mg/kg) starting 30 minutes after TBI and then every 24 hours for 7 days. RR behavioral assessment was conducted pre-TBI and on days 1-7, 14, 21, and 28 post-TBI. NSS was conducted pre-TBI and on days 1 and 28 post-TBI. Total improvement was determined by subtracting the day 1 score from the day 28 score.

### Immunofluorescence and Immunohistochemistry Analyses

Quantitative analyses of vascular area and neuronal density were performed by immunofluorescence microscopy using antibodies against CD31 and NeuN^7^ to label endothelial cells and neurons, respectively. Microglial and macrophage density was assessed by immunohistochemistry using an antibody against F4/80 ^17^. Vascular and neuronal densities were quantified in the striatum, whereas microglial/macrophage density was quantified in the cortex and hippocampus. For each region, analyses were performed in both the hemisphere ipsilateral to the lesion and the contralateral hemisphere for comparison.

## RESULTS

### rGDF11 promotes recovery post-ICH

Building on our previous work optimizing rGDF11 dosing after experimental ischemic stroke^5^, we hypothesized that treatment within the first week after ICH could similarly enhance recovery. A total of 22 animals were randomized to each group (vehicle and rGDF11). The first dose of rGDF11 was administered 30 minutes post-ICH, modeling early post-injury intervention, with daily dosing for 7 days, consistent with potential in-hospital therapy (Figure 1A). Over the 28-day study period, 7 animals in the vehicle group and 3 animals in the rGDF11 group died prior to the predefined study endpoint and were not included in behavioral or histological analyses. The study was not powered for survival, and no significant difference was observed between groups at 28 days (Figure 1B).

**Figure 1.**
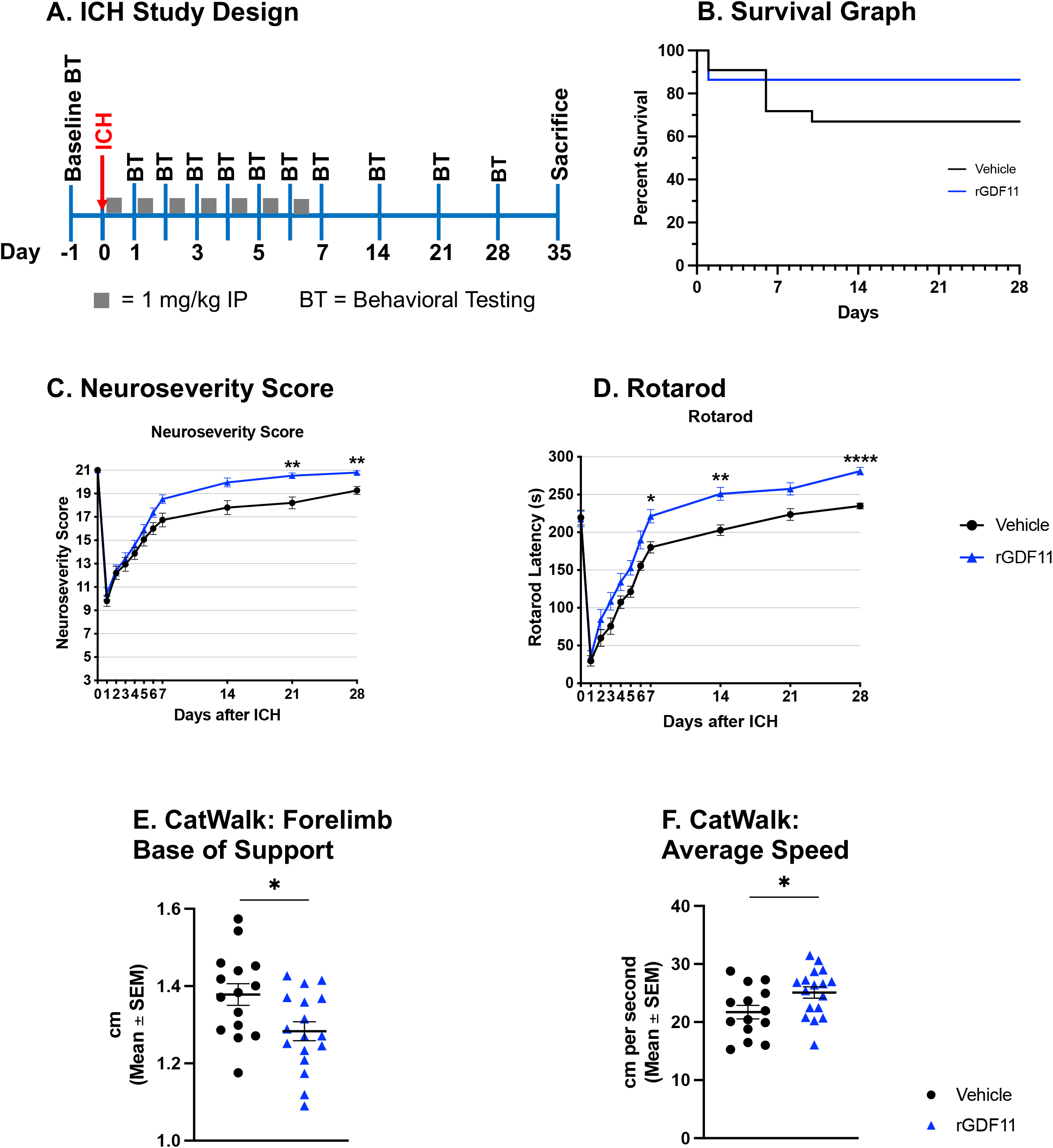
rGDF11 promotes sensorimotor recovery in a mouse model of intracerebral hemorrhage (ICH). **(A)** Schematic of the ICH study design indicating the day of ICH surgery, behavioral testing (BT) days, dosing days (gray squares), and day of termination. **(B)** Survival curve comparing vehicle- (n = 22) and rGDF11-treated (n = 22) mice. Log-rank (Mantel-Cox) test: *p* = 0.19. **(C)** Neuroseverity scores over time for vehicle- (n = 15) and rGDF11-treated (n = 19) mice. Two-way ANOVA (Geisser-Greenhouse correction): significant time x treatment interaction (*F* (10, 320) = 3.91, *p* < 0.0001). **(D)** Rotarod performance over time for vehicle- (n = 15) and rGDF11-treated (n = 19) mice. Two-way ANOVA (Geisser-Greenhouse correction): significant time x treatment interaction (*F* (10, 320) = 2.51, *p* = 0.0065). **(C-D)** Sidak’s multiple comparisons test was used for individual timepoint comparisons. **(E-F)** CatWalk gait analysis was performed on day 7 post-ICH, assessing **(E)** forelimb base of support (vehicle: n = 15; rGDF11: n = 17) and **(F)** average speed (vehicle: n = 14; rGDF11: n = 17). Data were analyzed using an unpaired *t*-test with Holm-Sidak correction. **(C-F)** Data are presented as mean ± SEM. Statistical significance is indicated as \**p* < 0.05; \*\**p* < 0.01; \*\**\*p* < 0.001; \*\*\*\**p* < 0.0001.

NSS scores were significantly improved in rGDF11-treated mice (n = 19) compared to vehicle-treated controls (n = 15) at 21 and 28 days (F(10, 320) = 3.91, p < 0.0001; Figure 1C). Notably, a trend toward early divergence in the first week was observed, consistent with our prior ischemic stroke studies ^5^. Performance on RR similarly demonstrated improved latency in rGDF11-treated mice at 7, 14, and 28 days, with early trends toward separation of treatment curves in the first week (F(10, 320) = 2.51, p = 0.0065; Figure 1D). Analysis of CW locomotor data at 7 days post-ICH revealed significant (unpaired *t*-test with Holm-Sidak correction; p<0.05) improvements in forelimb base of support and average speed in rGDF11-treated mice (Figure 1E-F). Pre-ICH assessments showed no differences between groups (data not shown). Overall, rGDF11 treatment during the first post-injury week led to improved sensorimotor function and gait performance after ICH.

### rGDF11 increases vascularization and neuronal density post-ICH

To evaluate the long-term histological effects of rGDF11 following ICH/TBI, brain tissue was collected 35 days post-injury (28 days after the final dose) and analyzed for markers of neuroinflammation, vascular remodeling, and neurogenesis. rGDF11 treatment significantly (p=0.0267) increased striatal CD31^+^ vascular area in the ipsilateral hemisphere and showed a trend toward elevation (p=0.0619) in the contralateral hemisphere (Figure 2A), consistent with previously described vascular effects of GDF11 ^5,18,19^. Furthermore, vascular junction density was significantly increased in both the ipsilateral (p=0.0473) and contralateral (p=0.0078) hemispheres (Figure 2B). A modest increase in mature neurons was also detected in the ipsilateral (p=0.053) and contralateral (p=0.0876) striatum (Figure 2C). rGDF11 treatment was associated with a significant reduction in F4/80^+^ cell density in in the ipsilateral cortex (p=0.039) and hippocampus bilaterally (ipsilateral: p=0.0359; contraletral: p=0.0360). (Figure 2D–E). Together, these findings suggest that rGDF11 may exert vascular, neuronal, and anti-inflammatory effects that persist beyond the treatment window, although additional mechanisms likely contribute to its early and long-term benefits after ICH.

**Figure 2.**
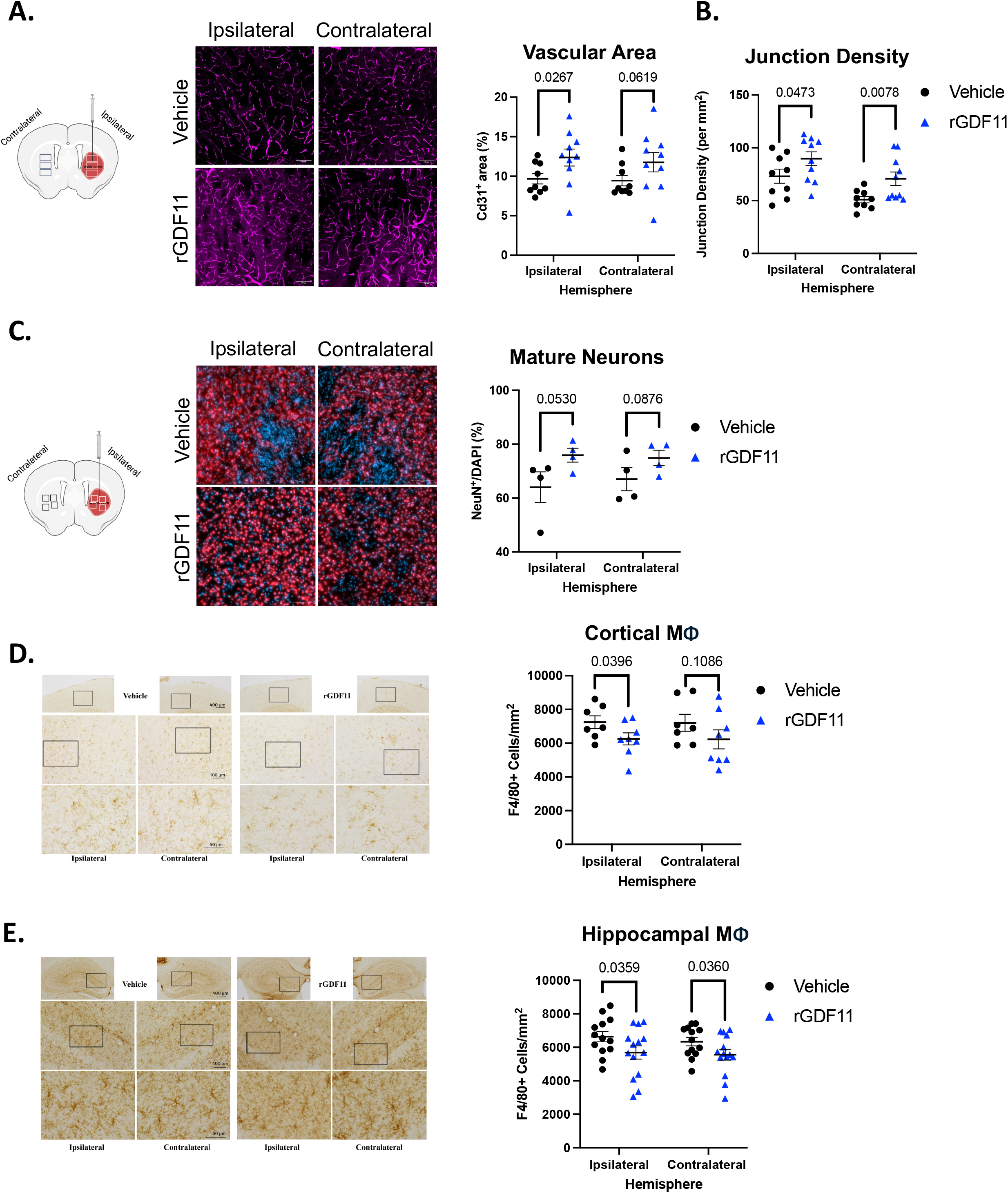
Histological analyses at 35 days post-ICH reveal increased vascularization and altered cellular composition in brains of rGDF11-treated animals. **(A)** Representative immunofluorescence images and quantification of CD31^+^ vascular area (% area covered) in the striatum (vehicle: *n* = 9; rGDF11: *n* = 10). **(B)** Quantification of vascular junction density in the striatum. **(C)** Representative immunofluorescence images and quantification of NeuN^+^ neurons as a percentage of total DAPI^+^ cells in the subventricular zone (vehicle: *n* = 4; rGDF11: *n* = 4). **(D-E)** Representative immunohistochemistry images and quantification of F4/80^+^ microglial/macrophage density in the cortex (**D**; vehicle: *n* = 7; rGDF11: *n* = 8) and hippocampus (**E**; vehicle: *n* = 13; rGDF11: *n* = 14). **(A-E)** All analyses were performed 35 days after ICH. Data are presented as mean ± SEM. Statistical comparisons were performed using one-tailed unpaired *t*-tests; *p*-values are shown for each comparison.

### rGDF11 promotes recovery post-TBI

Because results with rGDF11 following experimental ischemic stroke and ICH were remarkably similar, we decided to evaluate the benefits of rGDF11 after TBI. A total of 20 animals were randomized to each group (vehicle and rGDF11). We used the same rGDF11 treatment protocol as in ICH experiments, as TBI can present as an emergency, with the first dose 30 min after the trauma and then daily treatment for 7 days (Figure 3A). Over the 28-day study period, 8 animals in the vehicle group and 8 animals in the rGDF11 group died prior to the predefined study endpoint and were not included in behavioral analyses. There was no difference in survival between groups (Figure 3B).

**Figure 3.**
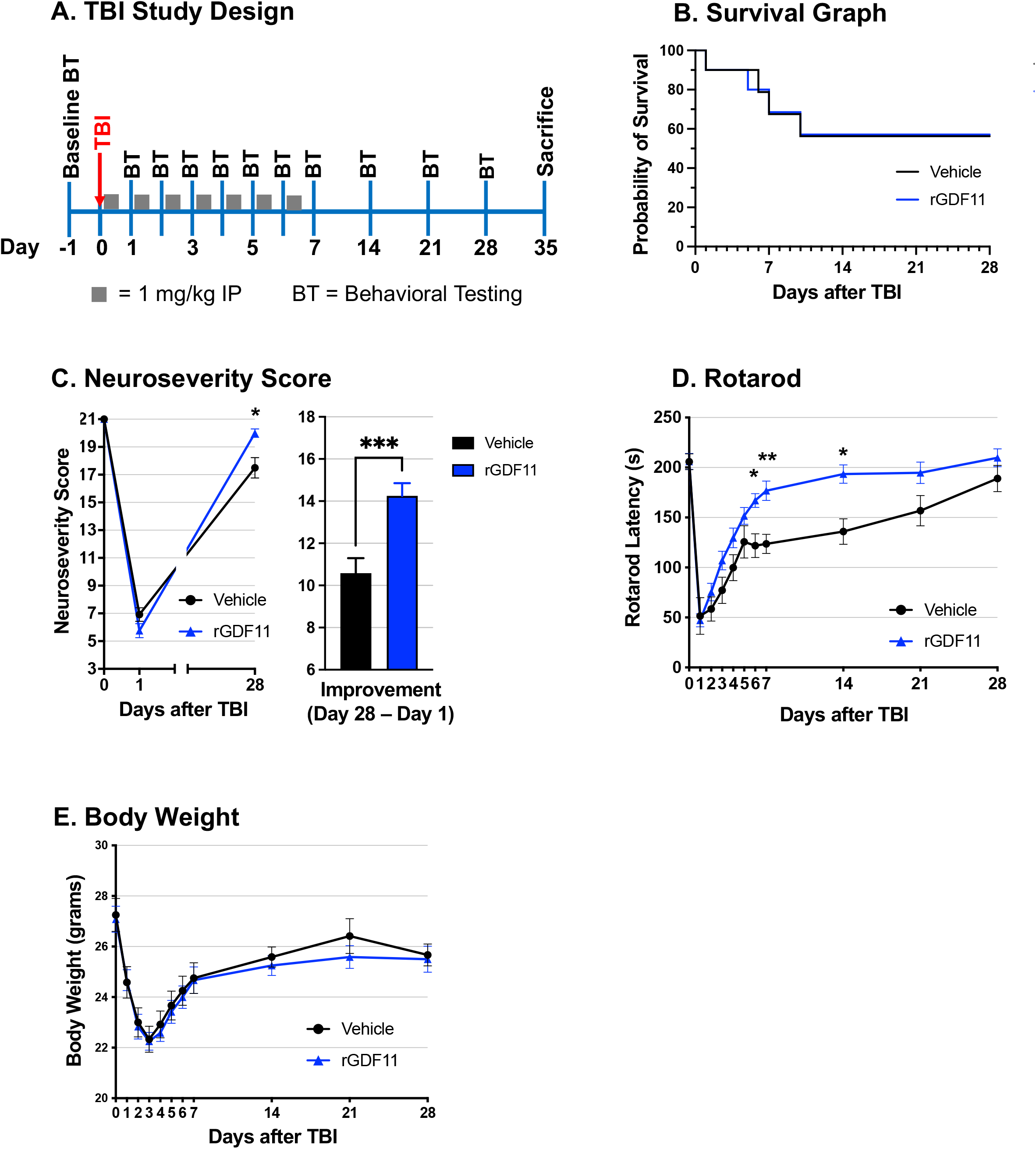
rGDF11 promotes sensorimotor recovery in a mouse model of traumatic brain injury (TBI). **(A)** Schematic of the TBI study design indicating the day of TBI (day 0), behavioral testing (BT) days, dosing days (gray squares), and day of termination. **(B)** Survival curve comparing vehicle- (n = 20) and rGDF11-treated (n = 20) mice. Log-rank (Mantel-Cox) test: *p* > 0.99. **(C)** Left panel: Neuroseverity scores pre-TBI, Day 1, and Day 28 for vehicle- (n = 12) and rGDF11-treated (n = 12) mice. Two-way ANOVA (Geisser-Greenhouse correction): significant time x treatment interaction (*F* (1.80, 39.60) = 10.55, *p* = 0.0003). Right panel: Neuroseverity score improvement on Day 28 compared to Day 1. Unpaired Welch’s t-test (two-tailed) **(D)** Rotarod performance over time for vehicle- (n = 12) and rGDF11-treated (n = 12) mice. Two-way ANOVA (Geisser-Greenhouse correction): significant time x treatment interaction (*F* (5.278, 116.1) = 2.53, *p* = 0.0303). **(E)** Body weight over time for vehicle- (n = 12) and rGDF11-treated (n = 12) mice. Two-way ANOVA (Geisser-Greenhouse correction): no significant time x treatment interaction (*F* (3.707, 81.56) = 0.211, *p* = 0.9214) **(C-E)** Sidak’s multiple comparisons test was used for individual timepoint comparisons. Data are presented as mean ± SEM. Statistical significance is indicated as \**p* < 0.05; \*\**p* < 0.01, \*\*\**p* < 0.001.

NSS scores were significantly improved in rGDF11-treated mice (n = 12) compared with vehicle-treated controls (n = 12) over time (F(1.803, 66.40) = 10.85, p < 0.001; Figure 3C, left panel). In addition, the magnitude of improvement from Day 1 to Day 28 was significantly greater in rGDF11-treated mice compared with vehicle-treated mice (p<0.001; Figure 3C, right panel).

RR performance for rGDF11-treated mice was significantly improved at 6, 7, and 14 days compared to vehicle-treated mice, although both groups were similar at 21 and 28 days (F(5.278, 116.1) = 2.53, p = 0.0303; Figure 3D). We also confirmed that rGDF11 treatment did not result in significant body weight loss following TBI, with no significant time × treatment interaction (F(3.707, 81.56) = 0.211, p = 0.9214; Figure 3E). These data suggest that rGDF11 may improve outcomes after TBI, although the optimal dosing to achieve sustained benefits will require further experimentation.

## DISCUSSION

ICH and TBI represent distinct forms of acute brain injury, one resulting from vascular rupture and bleeding, the other from mechanical impact and shearing forces. Despite these differences, both conditions share convergent downstream mechanisms, including vascular disruption, neuroinflammation, neuronal loss, and impaired tissue repair. These overlapping pathophysiological pathways highlight the need for neurorestorative approaches that target fundamental processes of brain recovery rather than injury-specific triggers. In this study, we demonstrate that recombinant GDF11 (rGDF11) enhances functional recovery in mouse models of both ICH and TBI, extending its previously reported benefits in ischemic stroke.^5-7^

A key feature of this work is that rGDF11 treatment was initiated after injury rather than as a pretreatment. This distinction is critical for translational relevance, as prophylactic paradigms rarely reflect clinical reality. Although treatment was initiated at 30 minutes post-injury, an early time point that may not be feasible in many real-world settings, this design provides important proof-of-concept that post-injury administration of rGDF11 can confer significant and durable benefit. While we did not systematically define the outer limits of the therapeutic window, these findings establish a foundation for future studies to determine whether efficacy can be extended to later and more clinically practical time points. Notably, functional improvements persisted for weeks following only a 7-day dosing regimen, suggesting that transient rGDF11 exposure may initiate sustained neurorestorative processes. These sustained effects are consistent with our prior findings in ischemic stroke, where transient rGDF11 treatment promoted long-term recovery.^5^

Across studies, a consistent observation has been the emergence of early functional improvement within the first week of rGDF11 administration. This reproducible trend across distinct injury paradigms, ischemic stroke ^5^, ICH, and TBI, suggests that rGDF11 rapidly engages endogenous repair pathways that transcend injury type. Although these early effects did not reach statistical significance in every experiment, their consistency supports further investigation into the acute molecular and cellular mechanisms underlying rGDF11’s actions. Future studies should focus on characterizing the temporal profile of rGDF11-mediated repair, including early biomarkers of vascular remodeling, neuroinflammation, and synaptic plasticity, to elucidate how rGDF11 initiates and sustains recovery following brain injury.

A limitation of the present study is the use of young adult male mice rather than aged or mixed-sex cohorts, which would more closely approximate the clinical population affected by ICH and TBI. This approach was selected to reduce variability related to age- and sex-dependent differences in neuroinflammatory and regenerative responses, thereby increasing sensitivity to detect treatment effects while minimizing the number of animals used. The encouraging functional improvements observed with rGDF11 treatment warrant future studies to evaluate dose-response relationships and treatment regimens across both sexes and in aged rodents. Histological analyses were performed on a limited sample size at 35 days post-ICH, and future studies will further investigate the molecular and cellular mechanisms underlying rGDF11-mediated neurorepair, including analyses at additional time points. Together, these findings provide a foundation for continued translational development of rGDF11 as a potential restorative therapy for ICH and TBI.

## ABBREVIATIONS

**Abbreviation Definition**

ARRIVE: Animal Research: Reporting of In Vivo Experiments
BSA: Bovine Serum Albumin
CD31: Cluster of Differentiation 31 (Endothelial cell marker)
CHO: Chinese Hamster Ovary
CW: CatWalk
DAPI: 4′,6-Diamidino-2-Phenylindole
F4/80: EGF-like module-containing mucin-like hormone receptor-like 1 (EMR1; Microglial marker)
GDF11: Growth Differentiation Factor 11
HEK: Human Embryonic Kidney
IACUC: Institutional Animal Care and Use Committee
ICH: Intracerebral Hemorrhage
MΦ: Macrophage and Microglia
NeuN: Neuronal Nuclei (neuronal marker) NIH National Institutes of Health
NSS: Neuroseverity Score
PBS: Phosphate-Buffered Saline
PBS-T: PBS with Tween-20
rGDF11: Recombinant Growth Differentiation Factor 11 ROI Region of Interest
RR: Rotarod Latency
TBI: Traumatic Brain Injury
TGF-β: Transforming Growth Factor-beta
ZFN: Zinc Finger Nuclease

## DATA AVAILABILITY

The data that support the findings of this study are available from the corresponding author upon reasonable request.

## ACKNOWLEDGMENTS

The authors have no acknowledgments to report.

## SOURCES OF FUNDING

Not applicable

## DISCLOSURES

Richard Lee, Manisha Sinha, Yongting Wang, Ori Cohen, and Anthony Sandrasagra are inventors on GDF11 patents. Richard Lee is on the Scientific Advisory Board of Alevian, Inc.

## REFERENCES

1. Magid-Bernstein J, Girard R, Polster S, Srinath A, Romanos S, Awad IA, Sansing LH. Cerebral Hemorrhage: Pathophysiology, Treatment, and Future Directions. Circ Res. 2022;130:1204–1229. doi: 10.1161/CIRCRESAHA.121.319949

2. Maas AIR, Menon DK, Manley GT, Abrams M, Akerlund C, Andelic N, Aries M, Bashford T, Bell MJ, Bodien YG, et al. Traumatic brain injury: progress and challenges in prevention, clinical care, and research. Lancet Neurol. 2022;21:1004–1060. doi: 10.1016/S1474-4422(22)00309-X

3. Ben Driss L, Lian J, Walker RG, Howard JA, Thompson TB, Rubin LL, Wagers AJ, Lee RT. GDF11 and aging biology - controversies resolved and pending. J Cardiovasc Aging. 2023;2023. doi: 10.20517/jca.2023.23

4. Loffredo FS, Steinhauser ML, Jay SM, Gannon J, Pancoast JR, Yalamanchi P, Sinha M, Dall’Osso C, Khong D, Shadrach JL, et al. Growth differentiation factor 11 is a circulating factor that reverses age-related cardiac hypertrophy. Cell. 2013;153:828–839. doi: 10.1016/j.cell.2013.04.015

5. Cohen OS, Sinha M, Wang Y, Daman T, Li PC, Deatherage C, Charrez B, Deshpande A, Jordan S, Makoni NJ, et al. Recombinant GDF11 Promotes Recovery in a Rat Permanent Ischemia Model of Subacute Stroke. Stroke. 2025;56:996–1009. doi: 10.1161/STROKEAHA.124.049908

6. Hudobenko J, Ganesh BP, Jiang J, Mohan EC, Lee S, Sheth S, Morales D, Zhu L, Kofler JK, Pautler RG, et al. Growth differentiation factor-11 supplementation improves survival and promotes recovery after ischemic stroke in aged mice. Aging (Albany NY). 2020;12:8049–8066. doi: 10.18632/aging.103122

7. Lu L, Bai X, Cao Y, Luo H, Yang X, Kang L, Shi M-J, Fan W, Zhao B-Q. Growth Differentiation Factor 11 Promotes Neurovascular Recovery After Stroke in Mice. Front Cell Neurosci. 2018;12:205. doi: papers3://publication/doi/10.3389/fncel.2018.00205

8. Percie du Sert N, Ahluwalia A, Alam S, Avey MT, Baker M, Browne WJ, Clark A, Cuthill IC, Dirnagl U, Emerson M, et al. Reporting animal research: Explanation and elaboration for the ARRIVE guidelines 2.0. PLoS Biol. 2020;18:e3000411. doi: 10.1371/journal.pbio.3000411

9. Wang H, Faw TD, Lin Y, Huang S, Venkatraman TN, Cantillana V, Lascola CD, James ML, Laskowitz DT. Neuroprotective Pentapeptide, CN-105, Improves Outcomes in Translational Models of Intracerebral Hemorrhage. Neurocrit Care. 2021;35:441–450. doi: 10.1007/s12028-020-01184-y

10. Lei B, James ML, Liu J, Zhou G, Venkatraman TN, Lascola CD, Acheson SK, Dubois LG, Laskowitz DT, Wang H. Neuroprotective pentapeptide CN-105 improves functional and histological outcomes in a murine model of intracerebral hemorrhage. Sci Rep. 2016;6:34834. doi: 10.1038/srep34834

11. Lei B, Sheng H, Wang H, Lascola CD, Warner DS, Laskowitz DT, James ML. Intrastriatal injection of autologous blood or clostridial collagenase as murine models of intracerebral hemorrhage. J Vis Exp. 2014. doi: 10.3791/51439

12. James ML, Sullivan PM, Lascola CD, Vitek MP, Laskowitz DT. Pharmacogenomic effects of apolipoprotein e on intracerebral hemorrhage. Stroke. 2009;40:632–639. doi: 10.1161/STROKEAHA.108.530402

13. Garcia JH, Wagner S, Liu KF, Hu XJ. Neurological deficit and extent of neuronal necrosis attributable to middle cerebral artery occlusion in rats. Statistical validation. Stroke. 1995;26:627–634; discussion 635. doi: 10.1161/01.str.26.4.627

14. Hamm RJ, Pike BR, O’Dell DM, Lyeth BG, Jenkins LW. The rotarod test: an evaluation of its effectiveness in assessing motor deficits following traumatic brain injury. J Neurotrauma. 1994;11:187–196. doi: 10.1089/neu.1994.11.187

15. Walter J, Kovalenko O, Younsi A, Grutza M, Unterberg A, Zweckberger K. The CatWalk XT(R) is a valid tool for objective assessment of motor function in the acute phase after controlled cortical impact in mice. Behav Brain Res. 2020;392:112680. doi: 10.1016/j.bbr.2020.112680

16. Laskowitz DT, Wang H, Chen T, Lubkin DT, Cantillana V, Tu TM, Kernagis D, Zhou G, Macy G, Kolls BJ, et al. Neuroprotective pentapeptide CN-105 is associated with reduced sterile inflammation and improved functional outcomes in a traumatic brain injury murine model. Sci Rep. 2017;7:46461. doi: 10.1038/srep46461

17. Lei B, Dawson HN, Roulhac-Wilson B, Wang H, Laskowitz DT, James ML. Tumor necrosis factor alpha antagonism improves neurological recovery in murine intracerebral hemorrhage. J Neuroinflammation. 2013;10:103. doi: 10.1186/1742-2094-10-103

18. Katsimpardi L, Litterman NK, Schein PA, Miller CM, Loffredo FS, Wojtkiewicz GR, Chen JW, Lee RT, Wagers AJ, Rubin LL. Vascular and neurogenic rejuvenation of the aging mouse brain by young systemic factors. Science. 2014;344:630–634. doi: papers3://publication/doi/10.1126/science.1251141

19. Ozek C, Krolewski RC, Buchanan SM, Rubin LL. Growth Differentiation Factor 11 treatment leads to neuronal and vascular improvements in the hippocampus of aged mice. Sci Rep. 2018;8:17293. doi: papers3://publication/doi/10.1038/s41598-018-35716-6

20. Laskowitz DT, Song P, Wang H, Mace B, Sullivan PM, Vitek MP, Dawson HN. Traumatic brain injury exacerbates neurodegenerative pathology: improvement with an apolipoprotein E-based therapeutic. J Neurotrauma. 2010;27:1983–1995. doi: 10.1089/neu.2010.1396

21. Zudaire E, Gambardella L, Kurcz C, Vermeren S. A computational tool for quantitative analysis of vascular networks. PLoS ONE. 2011;6:e27385. doi: 10.1371/journal.pone.0027385

