## Supplemental Methods for "Recombinant GDF11 Improves Neurological Recovery in Models of Hemorrhagic Stroke and Traumatic Brain Injury"

**Supplemental Material:**

**Supplemental Methods**

**Tables: N/A**

**Figure: N/A**

**References: 7, 8, 9, 10, 11, 12, 13, 14, 15, 16, 17, 20, 21**

### Supplemental Methods

#### Protein Production

Recombinant human GDF11 (rGDF11) mature dimer was produced through in-house upstream and downstream process development. Transient expression was first achieved in Expi293 cells using an rGDF11 expression construct to generate initial material and confirm protein expression. The same construct was subsequently used to develop a stable Chinese Hamster Ovary (CHO) cell line using the CHOZN® GS-/- ZFN-modified CHO platform (Millipore Sigma, Burlington, MA).

Briefly, the rGDF11 construct was introduced into CHOZN cells via electroporation, and electroporated populations were divided into minipools. These minipools were expanded and screened by ELISA to identify high-producing clones. Selected pools were further characterized by ELISA and western blot analysis to verify complete processing of the precursor protein into the mature GDF11 dimer. The top-performing minipool was subjected to single-cell cloning by limiting dilution, and individual clones were evaluated for both productivity and growth kinetics. The clone demonstrating optimal expression and doubling time was selected to generate research and master cell banks, which were subsequently established and characterized at Northway Biotech (Vilnius, Lithuania).

Purification of rGDF11 was performed in-house using a multistep chromatography workflow optimized for recovery of the mature dimeric form. The resulting protein was confirmed to be high-purity, properly processed, and biologically active. This material was used for all studies described in the present work.

**Animal Welfare Statement**

Male C57BL6J mice (12 – 14-week-old, The Jackson Laboratory, Bar Harbor, Maine, USA) were housed in a temperature and humidity-controlled environment with a 12 hour light-dark cycle and were allowed free access to food and water. The animals were randomized into experimental groups, all procedures, assessments, and data analyses were performed in a blinded fashion. All animal studies were approved by Duke University Institutional Animal Care and Use Committee and conducted in accordance with institutional and federal guidelines for animal care. Experiments adhered to ARRIVE Guidelines 2.0 to promote rigor, reproducibility, and transparency in animal research.[8]

**Intracerebral Hemorrhage (ICH) Model**

The intrastriatal collagenase model was performed as previously described[9,10,11], with minor modifications. Briefly, male C57BL/6J mice (12–14 weeks; Jackson Laboratories, Bar Harbor, ME) were anesthetized with 4.6% isoflurane for induction, followed by tracheal intubation and mechanical ventilation (tidal volume 0.3 mL; 140 breaths/min). Anesthesia was maintained with 1.6% isoflurane in a 30% $O_2$/70% $N_2$ gas mixture. Core temperature was maintained at 37 ± 0.2 °C using a rectal probe and circulating warm-water pad. The head was secured in a stereotactic frame, and a midline scalp incision was made to expose the skull. A burr hole was drilled lateral to bregma (2.2 mm). Type IV-S *Clostridium histolyticum* collagenase (Sigma-Aldrich) was injected over 5 min via a Hamilton syringe (Hamilton Company, Reno, NV) advanced to a depth of 3 mm (0.2 U in 0.32 µL saline). The needle was withdrawn slowly over 5 min, and bone wax was applied to seal the burr hole. The incision was closed with sutures, and animals were allowed to recover spontaneous respiration and righting reflex before extubating.

**Traumatic Brain Injury (TBI) Model**

The closed-head injury model was performed as previously described.[16] This model produces injury to selectively vulnerable neurons in the cortex and hippocampus and results in vestibulomotor and neurocognitive deficits. Male C57BL/6J mice (12–14 weeks; Jackson Laboratories, Bar Harbor, ME) were anesthetized with 4.6% isoflurane for induction and mechanically ventilated with 1.6% isoflurane in 30% $O_2$/70% $N_2$. Core temperature was maintained at 37 °C. To prevent basilar skull fracture, ear bars were not used. Mice were secured prone in a stereotactic frame on a molded acrylic cast, with surgical tape across the shoulders and the intubation tube secured to the cast. The cast provided 3 mm of space beneath the head to permit acceleration–deceleration during impact. After shaving the scalp and identifying bregma, a concave 3-mm metal disc was affixed to the skull just caudal to bregma. A 2-mm pneumatic impactor (Air-Power Inc., High Point, NC) delivered a single midline impact (velocity = 6.8 ± 0.2 m/s; displacement = 3 mm). Mice regained spontaneous ventilation prior to extubation and were then allowed free access to food and water. Body weights were recorded for all mice pre-TBI and on days 1–7, 14, 21, and 28 post-TBI.

**Drug Administration, Blinding, and Group Sizes**

For both ICH and TBI studies, recombinant GDF11 (rGDF11; 1 mg/kg) or vehicle was administered by intraperitoneal injection beginning 30 minutes post-injury and continued once daily for 7 days. Animals were randomly assigned to treatment groups, and investigators were blinded to treatment allocation throughout the study, including all behavioral and histological assessments. Treatment blinding and labeling were performed by an individual not involved in the animal procedures or data analysis, and group assignments were unblinded only after completion of all analyses. Group sizes were determined based on historical data from similar model studies to ensure adequate statistical power for behavioral outcomes while accounting for expected post-procedure mortality, resulting in 22 animals per treatment group for the ICH study[9,10,11] and 20 animals per treatment group for the TBI study[16,20].

**Behavioral Assessments**

Animals were acclimated to the behavioral testing environment for one week prior to baseline testing. All assessments were conducted under dim lighting and minimal background noise to reduce stress.

For ICH studies, behavioral testing included the Neuroseverity Score (NSS)[9,12,13], Rotarod (RR)[9,14], and CatWalk (CW)[15]. NSS and RR were measured pre-ICH and on days 1–7, 14, 21, and 28 post-ICH. CatWalk assessments were performed on day 7 post-injury.

For TBI studies, RR testing was performed pre-TBI and on days 1–7, 14, 21, and 28 post-TBI. NSS was evaluated pre-TBI and on days 1 and 28 post-injury. Functional improvement was determined by the change in score between day 1 and day 28.

**Rotarod Testing**

Vestibulomotor coordination was assessed an automated accelerating rotarod (RR) an automated accelerating rotarod (RR) (Rotarod ENV-577M, Med Associates Inc., Georgia, VT) as described previously.[9,14] Firstly, mice underwent a one-time acclimation for 300 second period in an acceleration speed of 2-20 rpm.  After the acclimation, the ability of the mice to remain on the accelerating RR was measured for three trials, each separated by 15 minutes to calculate an average score for each animal. Test run for a maximum of 300s with an accelerating rotational speed mode 4-40 rpm. Mice were tested on days 1 and 2 prior surgery to establish the baseline and on days 1-7, 14 and 28 post surgery. Baseline latencies were compared between groups to confirm equivalent pre-injury performance.

**Neuroseverity Score (NSS)**

Neurological function was assessed using a modified observational scoring system that evaluates seven domains: spontaneous activity, symmetry, climbing, balance/coordination, proprioception, vibrissae, and tactile responses.[9,12,13] Each domain was scored from 0 or 1–3 (minimum = 3, maximum = 21), with lower scores indicating greater impairment. All animals achieved the maximum score prior to injury. Average daily NSS values were used for group comparisons over time.

**CatWalk XT® Gait Analysis**

Gait parameters were assessed using the CatWalk XT system (Noldus Information Technology, Leesburg, VA) following established protocols.[15] Mice were acclimated to the testing room and apparatus, then trained to traverse the glass walkway continuously prior to baseline assessment. During testing, paw contacts on the internally illuminated glass plate were captured via high-speed video, in which paw placements appear bright against a dark body silhouette. All procedures were performed by an experimenter blinded to treatment group.

Locomotor data were collected from three successful runs at baseline and 7 days post-injury. A successful run was defined as uninterrupted ambulation across the walkway without stopping or sniffing; mice unable to complete the crossing within 5 seconds or those that died were excluded from analysis. Footprint detection and labeling were performed using CatWalk XT software (v10.6), followed by visual inspection and manual correction when necessary.

For each mouse, gait parameters, including run duration, average speed, forelimb and hindlimb base of support, cadence, regularity index, single-limb support, and variability measures, were averaged across the three valid runs and analyzed at 7 days post-injury. Outlier values exceeding two standard deviations from the group mean were excluded. Between-group comparisons were performed using independent-samples t-tests, with statistical significance set at $p < 0.05$.

**Perfusion and Histological Analyses**

Mice were anesthetized with isoflurane and intracardially perfused with saline at 35 days post-surgery. The removed brains were fixed in 4% buffered formaldehyde for 24 hours then transferred into 1X PBS at 4°C. Brains were incubated in 30% sucrose buffer for at least 24 hours, frozen coronal sections (40 μm) were cut and collected using a sliding microtome and stored in cryoprotectant solution containing 30% ethylene glycol, 15% sucrose and sodium phosphate.

**Vascular Analysis**

Vascular area was quantified by immunofluorescence microscopy using an antibody against CD31 to label endothelial cells.[7] Tissue sections were placed in 24-well plates containing PBS (MRGF6235, Growcells, Irvine, CA) and subjected to antigen retrieval with 1× Citrate Buffer (Ab93678, Abcam, Cambridge, MA) at 90°C for 10 min. Sections were then blocked with 10% Normal Donkey Serum (SD30-0500, VWR, Radnor, PA) in PBS containing 0.3% Triton X-100 (SLB2521, Sigma, Burlington, MA).

Sections were incubated overnight at 4°C on a shaking platform with primary antibody CD31 at 7.5 μg/mL (AF3628, R&D Systems, Minneapolis, MN) in 1% BSA in PBS, followed by three 5-min washes with PBS-T. The secondary antibody Donkey anti-Goat Alexa Fluor 647 (A32849, Thermo Fisher Scientific, Waltham, MA) was applied at 1:2500 dilution in PBS-T. Sections were counterstained with DAPI (62248, Thermo Fisher Scientific, Waltham, MA; 1:10,000 dilution), washed three times with PBS-T, and mounted on glass slides (22-035813, Thermo Fisher Scientific, Waltham, MA) using mounting medium (P36930, Thermo Fisher Scientific, Waltham, MA). Coverslips were sealed with nail lacquer (Amazon, WA).

Images were acquired from the ipsilateral striatum, in regions of interest (ROIs) surrounding the lesion, and from the contralateral striatum for comparison, using a Zeiss AxioScan Z1 slide-scanning microscope (ZEISS, Oberkochen, Germany) with a 20× objective.

Vascular area was quantified as the percentage of CD31-positive signal relative to total tissue area using Zeiss Zen Intellesis software (ZEISS, Oberkochen, Germany).

Vascular junction density was quantified with AngioTool software by identifying the number of junctions per ROI.[21] For each hemisphere, three ROIs were analyzed per section, and three coronal sections per brain, taken along the anterior–posterior axis at Bregma +1 mm, 0 mm, and –1 mm, were examined. Analyses included nine vehicle-treated and ten rGDF11-treated mice.

**Neuronal Analysis**

Neuronal density was quantified by immunofluorescence microscopy using an antibody against NeuN to label mature neurons.[7] Following the same antigen retrieval and blocking steps as described above, sections were incubated overnight at 4°C with primary antibody NeuN at 2 µg/mL (24307, Cell Signaling Technology, Danvers, MA) in 1% BSA in PBS. After washing, sections were incubated with secondary antibody Donkey anti-Rabbit Alexa Fluor 647 (A32795, Thermo Fisher Scientific, Waltham, MA) at 1:2500 dilution in PBS-T. Sections were counterstained with DAPI (1:10,000), washed, mounted, and coverslipped as described for vascular analysis.

Images were obtained from the **ipsilateral striatum** (region encompassing the lesion) and corresponding regions of the contralateral hemisphere using a Zeiss AxioScan Z1 slide-scanning microscope (ZEISS, Oberkochen, Germany) with a 20× objective. Neuronal density was expressed as the percentage of NeuN-positive cells relative to total DAPI-positive cells within each ROI, quantified using ImageJ software (NIH). Four ROIs were analyzed per hemisphere per coronal section at Bregma 0 mm. Analyses were performed on four brains per group for both vehicle- and rGDF11-treated mice.

#### Microglia and Macrophage Quantification

For microglia/macrophage staining with F4/80[17], equally spaced free-floating sections, 480 µm apart were incubated in 1% hydrogen peroxide, permeabilized by 0.1% saponin and blocked with 10% goat serum. Primary antibody (Rat anti-F4/80, 1:30,000; Invitrogen) was applied overnight at 4°C and then incubated in secondary antibody (biotinylated anti-rat IgG, 1:3000; Vector Laboratories) for 2 hours, followed by avidin-biotin-peroxidase complex treatment for 1 hour (ABC kit; Vector Laboratories). Staining was visualized with diaminobenzidine (DAB; Vector Laboratories). Between treatments, the sections were washed 3 times for 5 min each with Tris-buffered saline. The sections were then mounted onto charged slides and counterstained with hematoxylin (Fisher Scientific).

The quantification of F4/80-positive cells in hippocampus and medial cortex was done using the Optical Fractionator Probe, Stereo Investigator Software, MBF BioScience in all sections within location Bregma -1.06 mm to -2.46 mm. The stereology parameters for hippocampus and Cortex used were: Counting frame – 50 µm x 50 µm; Counting grid – 300 µm x 300 µm (hippocampus) or 400 µm x 400 µm (cortex); Section thickness – 13 µm; Dissector height – 10 µm. All the quantification was done double blinded.

#### Statistical Analyses

Statistical analyses were performed using Prism GraphPad (San Diego, CA). Animals that died prior to the predefined study endpoint were excluded from behavioral and histological analyses. Specific statistical tests and details for each analysis are provided in the corresponding figure legends. For behavioral and other analyses involving multiple comparisons, adjusted p-values are reported. For histological analyses, multiple testing correction was not applied given the limited number of planned comparisons. Statistical significance was defined as $p < 0.05$ unless otherwise noted.
